# Pharmacological Effects and Candidate Bioactive Constituents of Huanglian Wendan Decoction in Experimental Metabolic Syndrome: Multi-Omics Analysis of Gut–Liver Metabolism and NLRP3 Signaling

**DOI:** 10.64898/2026.10.05.756920

**Authors:** Hanyi Wang, Kaili Wang, Jincai Shi, Jianping Zhu

**Affiliations:** Hunan University of Chinese Medicine, 410208 Changsha, Hunan, China; Hunan Hospital of Integrated Traditional Chinese and Western Medicine, 410000 Changsha, Hunan, China

## Abstract

**Background:** Chronic low-grade inflammation and gut microbiota dysbiosis contribute to metabolic syndrome (MS). Huanglian Wendan Decoction (HLWDD) is a defined multi-herbal formulation used in Chinese medicine, but its effects on gut microbiota-associated metabolism and NLRP3 inflammasome signaling remain unclear.

**Objective:** This study evaluated the pharmacological effects and candidate absorbed bioactive constituents of HLWDD in experimental MS.

**Methods:** A high-fat, high-sugar diet-induced rat model of MS was assessed using metabolic, biochemical, histological, molecular, and multi-omics endpoints. Gut metagenomics, untargeted metabolomics of liver and colonic contents, liver transcriptomics, and UPLC-HRMS/MS analysis of serum-absorbed constituents were integrated with network pharmacology and molecular docking. Baicalin was further evaluated in PA-BSA-treated HepG2 cells.

**Results:** HLWDD ameliorated obesity, dyslipidemia, hepatic steatosis, and systemic inflammation and attenuated NLRP3 inflammasome-related signaling. Multi-omics analysis identified nine shared differential metabolites and a strong positive association between *Faecalibacterium* and butyrate. Thirty-two serum-absorbed constituents were tentatively identified. Baicalin showed favorable predicted interactions with NLRP3 (− 5.82 kcal/mol) and CASP1 ( − 5.41 kcal/mol) and concentration-dependently suppressed NLRP3 inflammasome-related signaling in HepG2 cells; the highest concentration increased intracellular butyrate.

**Conclusion:** HLWDD ameliorated metabolic and inflammatory abnormalities in experimental MS in association with restoration of gut–liver metabolic homeostasis and attenuation of NLRP3 inflammasome signaling. Baicalin emerged as a candidate absorbed bioactive constituent.

## Introduction

Metabolic syndrome (MS) is a prevalent metabolic disorder affecting more than one-quarter of the global population and is closely associated with obesity, type 2 diabetes, and non-alcoholic fatty liver disease [1–3]. Chronic low-grade inflammation is a major pathological feature of MS and contributes to dysregulated glucose and lipid metabolism, insulin resistance, and multi-organ dysfunction [4–6]. The NLRP3 inflammasome, an important sensor of metabolic danger signals, promotes caspase-1 activation and the maturation and release of interleukin-1β (IL-1β) and IL-18, thereby contributing to sustained metabolic inflammation [7,8]. Accumulating evidence implicates excessive NLRP3 activation in the development and progression of MS and its complications, making this pathway a potential target for metabolic-inflammatory regulation [9,10].

The gut microbiota interacts extensively with host metabolism through the gut–liver axis. In MS, gut dysbiosis is frequently accompanied by reductions in butyrate-producing bacteria and beneficial microbial metabolites, particularly short-chain fatty acids (SCFAs) [11–13]. Alterations in microbial composition and metabolite profiles may contribute to hepatic lipid accumulation and inflammatory responses, whereas restoration of microbial and metabolic homeostasis is associated with improved metabolic status [14,15]. However, the relationships linking specific microbial taxa and metabolites to host inflammatory pathways, including NLRP3 inflammasome signaling, remain incompletely understood.

Huanglian Wendan Decoction (HLWDD) is a defined multi-herbal formulation used in Chinese medicine. Previous experimental studies have reported that HLWDD improves dyslipidemia, insulin resistance, hepatic inflammation, and other metabolic abnormalities [16–18]. Additional evidence suggests that HLWDD may modulate gut microbial composition and microbial metabolites while attenuating inflammatory signaling, including NLRP3-related pathways [19,20]. These findings provide a pharmacological basis for investigating HLWDD as a multi-component intervention in experimental MS. However, how its effects on gut microbiota-associated metabolism relate to NLRP3 inflammasome signaling, and which absorbed constituents may contribute to these effects, remain unclear.

Therefore, we hypothesized that HLWDD ameliorates experimental MS through coordinated modulation of gut microbiota-associated metabolic profiles and NLRP3 inflammasome-related signaling. To test this hypothesis, a high-fat, high-sugar diet-induced rat model of MS was evaluated using objective metabolic, biochemical, histological, molecular, and multi-omics endpoints. Gut metagenomics, untargeted metabolomics of liver and colonic contents, and liver transcriptome sequencing were integrated with UPLC-HRMS/MS analysis of serum-absorbed constituents, network pharmacology, and molecular docking. Baicalin, identified as a candidate absorbed constituent, was further evaluated in a PA-BSA-induced HepG2 lipotoxicity model. No traditional Chinese medicine syndrome or pattern classification was used for animal selection, grouping, treatment allocation, or outcome assessment. This study aimed to characterize the pharmacological effects, associated gut – liver metabolic alterations, and candidate absorbed bioactive constituents of HLWDD in experimental MS.

## Materials and Methods

### Reagents and Antibodies

Metformin (batch no. 20200A; Patheon France, France) was used as the positive control. RNA extraction (NovoStart®, batch no. 5003050), cDNA synthesis (NovoScript®, batch no. E047-01B), and SYBR Green qPCR (NovoStart®, batch no. E099-01A) kits were purchased from Novoprotein Scientific Inc. Serum biochemical assay kits for total cholesterol (TC, batch no. M130055-48T), triglycerides (TG, batch no. BES2945K), high-density lipoprotein cholesterol (HDL-C, batch no. BES5270K), and low-density lipoprotein cholesterol (LDL-C, batch no. BES5287K) were obtained from Biosharp Life Sciences.

ELISA kits for rat and human TNF-α, IL-1 β, IL-6, IL-4, IL-10, and IL-13 were purchased from Shenzhen Xinbosheng Biotechnology Co., Ltd. CCK-8 reagent was obtained from Wuhan Elabscience Biotechnology Co., Ltd. The following primary antibodies were used: NLRP3 (ab263899) and IL-1β (ab283818) from Abcam; ASC (A22046), GSDMD (A20197), and caspase-1 (A25308) from ABclonal; and IL-18 (10663-1-AP) from Proteintech.

### Preparation of HLWDD

The herbal ingredients were weighed according to the prescribed composition: *Coptis chinensis* Franch. (rhizome; Sichuan, China; batch no. CC202305), 10 g; *Pinellia ternata* (Thunb.) Makino (prepared tuber; Shanxi, China; batch no. PT202304), 10 g; *Phyllostachys edulis* (Carrière) J.Houz. (caulis; Zhejiang, China; batch no. PE202306), 10 g; *Citrus aurantium* L. (immature fruit; Jiangxi, China; batch no. CA202302), 10 g; *Citrus reticulata* Blanco (pericarp; Guangdong, China; batch no. CR202303), 10 g; *Glycyrrhiza uralensis* Fisch. ex DC. (honey-processed root; Inner Mongolia, China; batch no. GU202307), 5 g; *Zingiber officinale* Roscoe (fresh rhizome; Shandong, China; batch no. ZO202305), 5 g; and *Wolfiporia cocos* (F.A.Wolf) Ryvarden & Gilb. (sclerotium; Yunnan, China; batch no. WC202308), 15 g. The materials were morphologically authenticated, and botanical nomenclature was verified using the Medicinal Plant Names Service (MPNS). Voucher specimens were deposited in the institutional herbarium (accession nos. HLWD2023-001–008).

The herbs were decocted with eight volumes of water for 1 h and filtered, followed by a second decoction with six volumes of water for 30 min. The combined filtrates were concentrated, and the dosing solutions were subsequently adjusted to crude-drug equivalent concentrations of 0.36, 0.72, and 1.44 g/mL for the HLWDD-L, HLWDD-M, and HLWDD-H groups, respectively. For chemical analysis, a 10 mL aliquot was adjusted to 70% ethanol, refrigerated for 12 h, and filtered. Two milliliters of the filtrate was diluted with methanol, mixed with the internal standard solution, passed through a 0.45 μ m membrane, and centrifuged at 15,000 rpm for 5 min before analysis.

### Experimental Animals and Metabolic Syndrome Model Establishment

Forty-eight specific pathogen-free male Sprague–Dawley rats (200 ± 20 g; license no. SCXK [Xiang] 2025-0007) were housed at the Animal Center of Hunan University of Chinese Medicine under controlled conditions (20 ± 2 ° C, 40%–60% relative humidity, and a 12 h light/dark cycle). All procedures were approved by the Animal Ethics Committee of Hunan University of Chinese Medicine (LLBH-LL2025121708) and conducted in accordance with institutional and national guidelines for the care and use of laboratory animals.

After one week of acclimatization, eight rats were randomly assigned to the control group and fed a standard diet, whereas the remaining 40 rats received a high-fat, high-sugar diet containing 67% basal diet, 20% sucrose, 10% lard, 2% cholesterol, and 1% bile salts for 20 weeks to induce experimental MS. Model establishment was evaluated exclusively on the basis of objective metabolic and biochemical characteristics, including changes in body weight, fasting blood glucose, and serum lipid-related parameters relative to the control group.

### Grouping, Drug Administration, and Sample Collection

After successful establishment of the experimental MS model, the MS rats were randomly divided into five groups (n = 8 per group): model, metformin (400 mg/kg/day), HLWDD low-dose (HLWDD-L, 3.6 g/kg/day), medium-dose (HLWDD-M, 7.2 g/kg/day), and high-dose (HLWDD-H, 14.4 g/kg/day) groups [19]. The original control group (n = 8) was maintained throughout the treatment period. Group allocation and treatment evaluation were based exclusively on the experimental design and objective biomedical endpoints. While maintained on their respective diets, all groups received once-daily oral gavage for four consecutive weeks. The HLWDD-L, HLWDD-M, and HLWDD-H dosing solutions were prepared at crude-drug equivalent concentrations of 0.36, 0.72, and 1.44 g/mL, respectively, and administered at 10 mL/kg. Metformin was dissolved in distilled water at 40 mg/mL and administered at 10 mL/kg, corresponding to 400 mg/kg/day [21]. The control and model groups received an equivalent volume of distilled water.

After the final administration, rats were fasted overnight and anesthetized. Blood was collected from the abdominal aorta, and serum was separated by centrifugation at 3,000 rpm for 15 min at 4 °C. The rats were then euthanized, and liver tissues were rapidly collected, with portions snap-frozen in liquid nitrogen and the remainder fixed in 4% paraformaldehyde. Colonic contents were collected and stored at −80 °C until analysis.

### Pharmacodynamic Assessments

Serum total cholesterol (TC), triglycerides (TG), high-density lipoprotein cholesterol (HDL-C), low-density lipoprotein cholesterol (LDL-C), and fasting blood glucose (FBG) were measured using an automatic biochemical analyzer. Serum TNF-α, IL-1β, IL-6, IL-4, IL-10, and IL-13 levels were determined using ELISA kits according to the manufacturers’ instructions. Liver tissues fixed in 4% paraformaldehyde were paraffin-embedded, sectioned at 5 μ m, and stained with hematoxylin and eosin (H&E) for histopathological examination under a light microscope.

### Liver Transcriptome Sequencing

Total RNA was extracted from liver tissues using TRIzol reagent, and RNA integrity was assessed using an Agilent 2100 Bioanalyzer. Transcriptome libraries were constructed and sequenced on an Illumina NovaSeq platform. Differentially expressed genes between the model and HLWDD-H groups were identified using |log2FC| > 1 and FDR < 0.05, followed by Gene Ontology (GO) and Kyoto Encyclopedia of Genes and Genomes (KEGG) enrichment analyses.

### Quantitative Real-Time PCR

Total RNA was extracted from liver tissues and cultured cells using TRIzol reagent, and RNA purity was assessed by spectrophotometry. Following reverse transcription, quantitative real-time PCR was performed using β-actin or GAPDH as the reference gene, as appropriate. Target genes included *Nlrp3*, *Pycard*, *Casp1*, *Gsdmd*, *Il1b*, and *Il18* in rat liver tissues and *NLRP3*, *PYCARD*, *CASP1*, *GSDMD*, *IL1B*, and *IL18* in HepG2 cells. Relative mRNA expression was calculated using the 2^−ΔΔCt^ method.

### Western Blot Analysis

Total protein was extracted from liver tissues or cultured cells using RIPA lysis buffer containing protease inhibitors, and protein concentrations were determined using the BCA assay. Proteins were separated by SDS-PAGE and transferred onto PVDF membranes. After blocking with 5% BSA for 1 h at room temperature, the membranes were incubated overnight at 4 °C with primary antibodies against NLRP3, ASC, caspase-1, GSDMD, IL-1β, IL-18, and the corresponding loading controls. After washing, the membranes were incubated with HRP-conjugated secondary antibodies for 1 h at room temperature. Protein bands were visualized using ECL and quantified using ImageJ. The levels of NLRP3, ASC, cleaved CASP1, N-GSDMD, IL-1 β, and IL-18 were normalized to the corresponding loading control.

### Gut Metagenomic Sequencing

DNA was extracted from colonic contents using the PF Mag-Bind Stool DNA Kit. DNA concentration and purity were assessed, and integrity was evaluated by 1% agarose gel electrophoresis. DNA was fragmented using a Covaris M220 system, and fragments of approximately 400 bp were selected for paired-end library construction. Sequence reads were assembled using MEGAHIT, and contigs ≥ 300 bp were retained. Open reading frames (ORFs) were predicted using Prodigal, and genes ≥ 100 bp were translated into amino acid sequences. High-quality reads from each sample were mapped to the non-redundant gene catalog at 95% identity using SOAPaligner to calculate gene abundance profiles.

### Untargeted Metabolomics Analysis

Liver tissues and colonic contents were subjected to untargeted metabolomics using a Thermo Dionex UltiMate 3000 RSLC system coupled to a Thermo Q Exactive mass spectrometer. Chromatographic separation was performed on an HSS T3 column (100 mm × 2.1 mm, 1.8 μm). Mobile phase A consisted of 95% water and 5% acetonitrile containing 0.1% formic acid, and mobile phase B consisted of 47.5% acetonitrile, 47.5% isopropanol, and 5% water containing 0.1% formic acid. The positive-ion gradient was: 0–3 min, 0%–20% B; 3–4.5 min, 20% –35% B; 4.5–5 min, 35%–100% B; 5–6.3 min, 100% B; 6.3–6.4 min, 100%–0% B; and 6.4–8 min, 0% B. The negative-ion gradient was: 0–1.5 min, 0%–5% B; 1.5–2 min, 5%–10% B; 2–4.5 min, 10%–30% B; 4.5–5 min, 30%–100% B; 5–6.3 min, 100% B; 6.3–6.4 min, 100%–0% B; and 6.4–8 min, 0% B. The flow rate was 0.40 mL/min, and the column temperature was maintained at 40 °C. Mass spectra were acquired in both positive-and negative-ion modes. After data preprocessing, multivariate analyses were performed to identify differential metabolites.

### UPLC-HRMS/MS Analysis of Serum-Absorbed HLWDD Constituents

For LC-MS/MS analysis, 50 μ L of serum was mixed with 150 μ L of acetonitrile for protein precipitation, vortexed, and centrifuged at 13,000 × g for 10 min. A 2 μL aliquot of the supernatant was injected into a Thermo Dionex UltiMate 3000 RSLC system equipped with an ACQUITY UPLC BEH C18 column (2.1 × 100 mm, 1.7 μ m). The mobile phases were water containing 0.1% formic acid (A) and acetonitrile (B). The gradient was: 0–1 min, 5% B; 1–15 min, 5%–95% B; 15–18 min, 95% B; 18–23 min, 95%–5% B; and 23–30 min, 5% B for re-equilibration. The flow rate was 0.3 mL/min. Mass spectrometric analysis was performed using a Thermo Q Exactive mass spectrometer equipped with an electrospray ionization source in both positive-and negative-ion modes.

### Network Pharmacology and Molecular Docking

Potential targets of HLWDD constituents were predicted using the TCMSP and SwissTargetPrediction databases, and target names were standardized to official gene symbols using UniProt. MS-related genes were retrieved from GeneCards using “metabolic syndrome” as the keyword. Overlapping targets between HLWDD constituents and MS were identified using Venny 2.1.0. A protein – protein interaction (PPI) network was constructed using STRING, and core targets were screened using the CytoNCA plug-in in Cytoscape 3.9.0. The compound–target– pathway network was also constructed using Cytoscape 3.9.0. GO and KEGG enrichment analyses were performed using DAVID 6.8. Representative compounds and targets were selected for molecular docking using AutoDock 4.2.6, and docking poses were visualized using PyMOL 2.4.1. Ligand and receptor structures were obtained from PubChem and the Protein Data Bank (PDB), respectively.

### Determination of Baicalin in HLWDD-Containing Rat Serum by HPLC

An appropriate amount of baicalin reference standard was accurately weighed, dissolved in methanol, and diluted to prepare a stock solution. Blank rat serum was spiked with the baicalin stock solution to obtain a series of calibration standards ranging from 6.10 to 6,250.00 ng/mL. Each serum sample (100 μL) was mixed with 300 μL of methanol for protein precipitation, vortexed for 3 min, and centrifuged at 13,000 rpm for 10 min at 4 °C. The supernatant was filtered through a 0.22 μ m membrane before analysis. The calibration curve was constructed using weighted (1/x) linear regression of baicalin peak area (Y) against serum concentration (X, ng/mL).

Chromatographic separation was performed on an Elite C18 column (4.6 × 250 mm, 5 μm). The mobile phase consisted of methanol (A) and 0.2% phosphoric acid aqueous solution (B) at a ratio of 47:53 (v/v) under isocratic elution. The detection wavelength was 280 nm, with a flow rate of 1.0 mL/min, column temperature of 30 °C, and injection volume of 20 μL.

HLWDD-containing rat serum samples collected as described in the Grouping, Drug Administration, and Sample Collection section were processed using the same protein precipitation, centrifugation, and filtration procedures described above for the calibration standards. The resulting filtrates were subjected to HPLC analysis, and baicalin concentrations were calculated using the corresponding calibration curve.

### Cell Culture

Human hepatocellular carcinoma HepG2 cells obtained from the Cell Bank of the Chinese Academy of Sciences were cultured in William’s Medium E supplemented with 5% fetal bovine serum, 2 mM L-glutamine, 0.05 μg/mL hydrocortisone, 5 μg/mL insulin, and 10 mM HEPES at 37 °C in a humidified atmosphere containing 5% CO₂. Cells in the logarithmic growth phase were used for subsequent experiments.

### Cell Viability Assay and Lipotoxicity Model

HepG2 cells in the logarithmic growth phase were seeded in 96-well plates at a density of 5 × 10³ cells/well. After 24 h, the medium was replaced with fresh medium containing baicalin at 0, 10, 20, 40, 60, 80, or 100 μM, and cells were incubated for an additional 24 h. Cell viability was assessed using the CCK-8 assay by adding 10 μL of reagent per well, followed by incubation for 2 h and measurement of absorbance at 450 nm.

The palmitic acid–bovine serum albumin (PA-BSA) complex was prepared by dissolving 10 mM palmitic acid in 0.1 M NaOH at 75 °C and mixing it with 3.33 mM BSA at 50 °C for 1 h to obtain a 5 mM PA-BSA solution (molar ratio 3:1, pH 7.4). The solution was filtered through a 0.22 μm membrane before use.

Cells were assigned to the control, model (0.3 mM PA-BSA), metformin (0.3 mM PA-BSA + 1.0 mM metformin) [22], and baicalin low-, medium-, and high-concentration groups (0.3 mM PA-BSA + 15, 30, or 60 μM baicalin, respectively). After 24 h of treatment, culture supernatants were collected, and TNF-α, IL-1β, IL-6, IL-4, IL-10, and IL-13 levels were determined using ELISA kits according to the manufacturers’ instructions.

### Detection of Gene and Protein Expression in Cells

Cell grouping and treatment were performed as described in the Cell Viability Assay and Lipotoxicity Model section. After treatment, cells were harvested for RNA and protein extraction according to the procedures described in the Quantitative Real-Time PCR and Western Blot Analysis sections. The mRNA levels of *NLRP3*, *PYCARD*, *CASP1*, *GSDMD*, *IL1B*, and *IL18* were determined by qRT-PCR. Protein levels of NLRP3, ASC, cleaved CASP1, N-GSDMD, IL-1 β, and IL-18 were determined by Western blotting, with GAPDH used as the corresponding reference control.

### Targeted Metabolomic Analysis of Intracellular Short-Chain Fatty Acids in HepG2 Cells

After treatment, cells were collected, washed twice with PBS, and lysed in 1 mL of pre-cooled methanol–water (1:1, v/v) by ultrasonication in an ice bath (200 W, 5 s on/5 s off for 5 min). The lysates were centrifuged at 14,000 rpm for 10 min at 4 °C, and the supernatants were collected. 2-Ethylbutyric acid was added as an internal standard at a final concentration of 100 μ M. After vortexing, an equal volume of diethyl ether was added for extraction, followed by vortexing for 3 min and centrifugation at 5,000 rpm for 5 min at 4 °C. The upper organic phase was collected, dried under nitrogen, and reconstituted in 50 μL of acetonitrile.

SCFAs were analyzed using gas chromatography–mass spectrometry with a DB-FFAP capillary column (30 m × 0.25 mm, 0.25 μm). Helium was used as the carrier gas at 1.0 mL/min. The oven temperature was maintained at 50 °C for 2 min, increased to 220 °C at 10 °C/min, and held for 5 min. The injector temperature was 250 °C with a split ratio of 10:1. Electron ionization was performed at 70 eV, with ion source and quadrupole temperatures of 230 and 150 °C, respectively. Acetate, propionate, butyrate, valerate, and caproate were quantified in selected-ion monitoring mode using the internal standard method.

### Statistical Analysis

Statistical analyses were performed using GraphPad Prism 9. Quantitative data are presented as the mean ± standard error of the mean (SEM), unless otherwise specified, and the number of independent biological replicates (n) is indicated in the corresponding figure legends. For datasets with an adequate sample size, normality and homogeneity of variance were assessed using the Shapiro – Wilk and Levene’s tests, respectively. Normally distributed data with homogeneous variances were analyzed by one-way ANOVA followed by the least significant difference (LSD) test. Normally distributed data with unequal variances were analyzed by Welch’s ANOVA followed by the Games– Howell test, whereas non-normally distributed data were analyzed using the Kruskal–Wallis test followed by Dunn’s multiple-comparison test.

Longitudinal measurements, including body weight, were analyzed using repeated-measures ANOVA followed by appropriate multiple-comparison tests. Pearson’s correlation analysis was used when parametric assumptions were satisfied, whereas Spearman’s rank correlation was used otherwise. For Western blotting and qRT-PCR experiments with n = 3, the replicates represented independent biological rather than technical replicates. All tests were two-tailed, and *P* < 0.05 was considered statistically significant.

## Results

### *In Vivo* Animal Studies

#### Therapeutic Effects of HLWDD on MS Rats

As shown in Figure 1A, body weight was markedly increased in the model group compared with the control group during the 20-week induction period. After four weeks of treatment, body weight and abdominal circumference were significantly reduced in the HLWDD-M and HLWDD-H groups compared with the model group (*P* < 0.05; Figure 1B,C).

**Fig 1.**
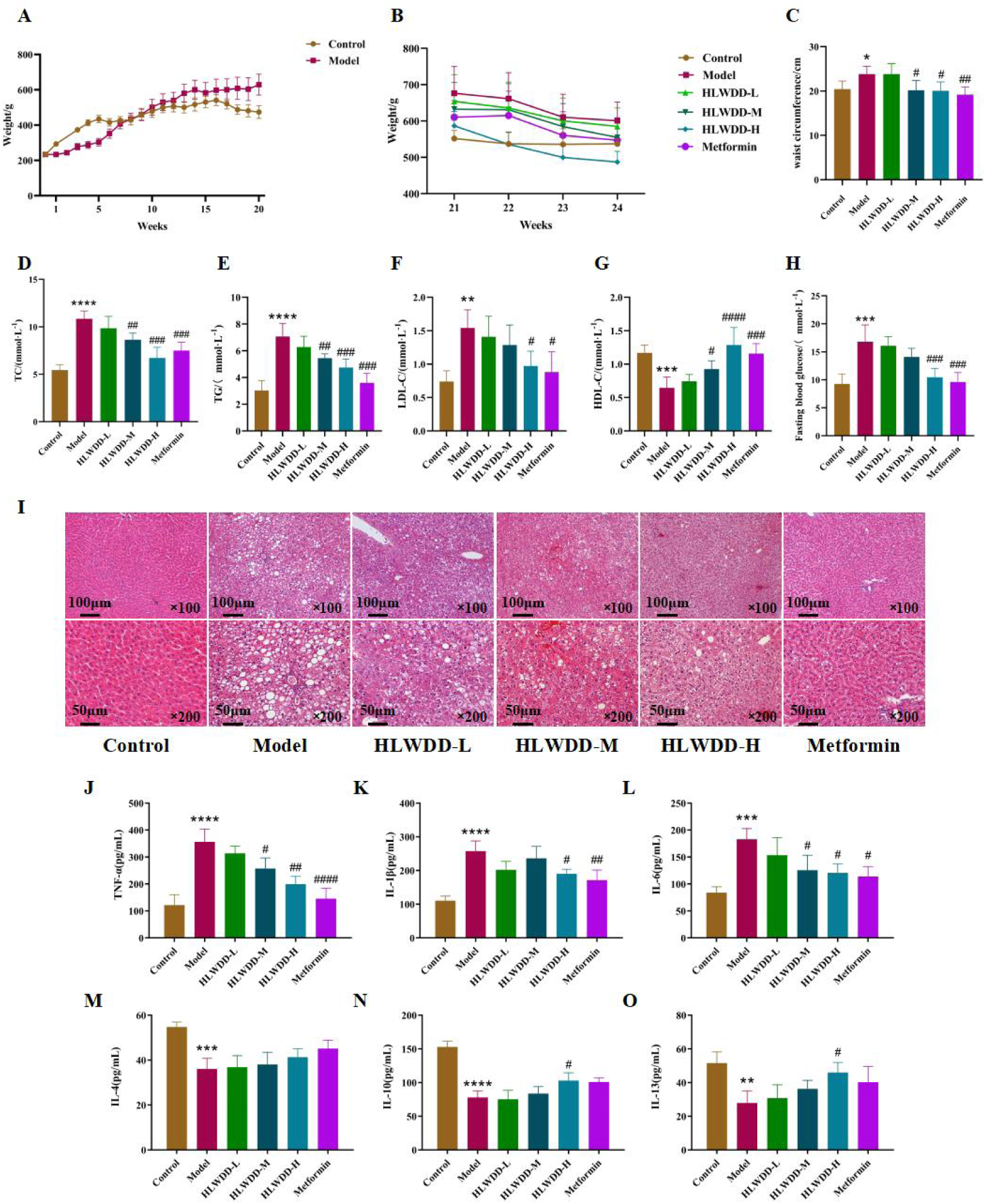
Therapeutic effects of HLWDD in MS rats. (A) Body weight changes during the 20-week model induction period; (B,C) body weight (B) and abdominal circumference (C) after four weeks of treatment; (D – H) serum metabolic parameters: total cholesterol (TC, D), triglycerides (TG, E), low-density lipoprotein cholesterol (LDL-C, F), high-density lipoprotein cholesterol (HDL-C, G), and fasting blood glucose (FBG, H); (I) representative H&E-stained liver sections at 100× and 200× magnification. Scale bars: 100 μm at 100× and 50 μm at 200×; (J–O) serum inflammatory cytokines: TNF-α (J), IL-1β (K), IL-6 (L), IL-4 (M), IL-10 (N), and IL-13 (O). Data are expressed as mean ± SEM (n = 3 biological replicates). \**P*<0.05, \*\**P*<0.01, \*\*\**P*<0.001, \*\*\*\**P*<0.0001, versus the control group; #*P*<0.05, ##*P*<0.01, ###*P*<0.001, ####*P*<0.0001, versus the model group.

Serum analysis showed that the model group exhibited increased TC, TG, LDL-C, and FBG (*P* < 0.01) and decreased HDL-C (*P* < 0.001) compared with the control group. HLWDD-M significantly reduced TC and TG (*P* < 0.01) and increased HDL-C (*P* < 0.05), whereas HLWDD-H reduced TC, TG, LDL-C, and FBG (*P* < 0.05) and increased HDL-C (*P* < 0.0001) (Figure 1D–H).

H&E staining showed normal hepatic architecture in the control group and pronounced steatosis in the model group. HLWDD treatment, particularly at the high dose, attenuated hepatic lipid accumulation and inflammatory infiltration, although mild hepatocyte edema remained (Figure 1I).

Compared with the control group, the model group showed increased serum TNF-α, IL-1β, and IL-6 (*P* < 0.001) and decreased IL-4, IL-10, and IL-13 (*P* < 0.01). HLWDD-M reduced TNF-α and IL-6 (*P* < 0.05), whereas HLWDD-H reduced TNF-α, IL-1β, and IL-6 (*P* < 0.05) and increased IL-10 and IL-13 (*P* < 0.05) (Figure 1J–O).

Collectively, these results indicate that HLWDD ameliorates metabolic abnormalities, hepatic steatosis, and systemic inflammation in MS rats.

#### Transcriptomic Analysis of the NLRP3 Inflammasome Pathway in Rat Liver

RNA-seq analysis showed that, compared with the model group, Nlrp3, Pycard, Casp1, and Il1b were significantly downregulated in the HLWDD-H group (log2FC < − 1, FDR < 0.05) (Figure 2A,B). Functional enrichment analysis indicated that the differentially expressed genes were mainly associated with inflammatory and innate immune pathways, including TNF, NOD-like receptor, Toll-like receptor, and MAPK signaling (Figure 2C), suggesting that NLRP3 inflammasome-related signaling may be involved in the effects of HLWDD.

**Fig 2.**
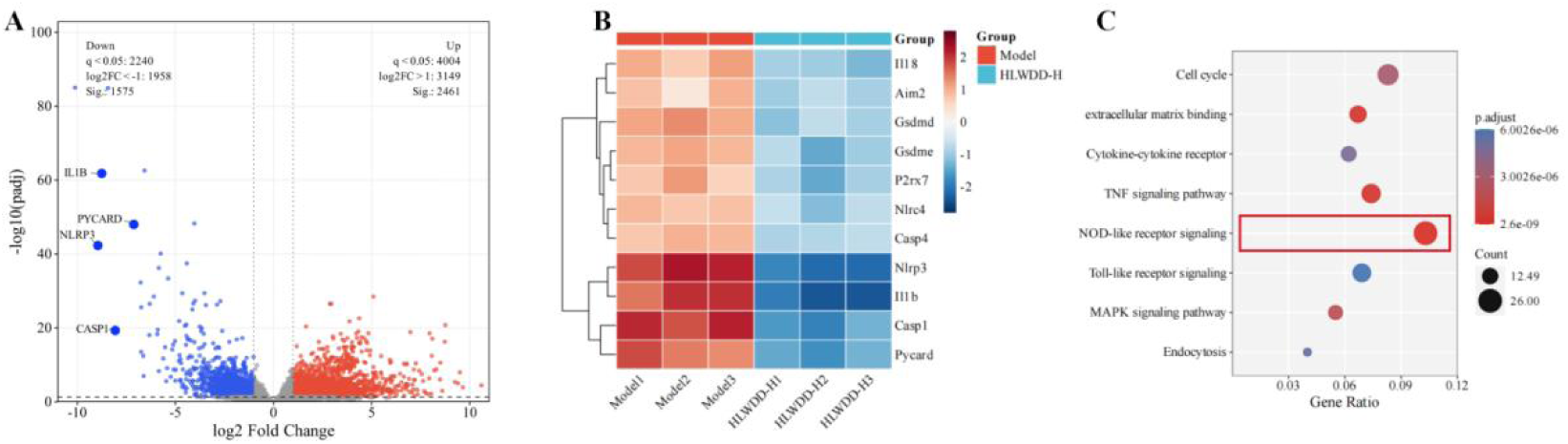
Transcriptomic analysis of liver tissues from the model and HLWDD-H groups. (A) Volcano plot of differentially expressed genes; (B) heatmap of representative differentially expressed genes; (C) functional enrichment analysis of differentially expressed genes.

#### Effects of HLWDD on the NLRP3 Inflammasome Pathway in Rat Liver

Western blot analysis showed that NLRP3, ASC, cleaved CASP1, N-GSDMD, IL-1 β, and IL-18 were significantly increased in the model group compared with the control group (*P* < 0.01), whereas HLWDD-M and HLWDD-H significantly reduced these protein levels (*P* < 0.05). At the mRNA level, HLWDD-H significantly reduced Nlrp3, Pycard, Casp1, Il1b, and Il18, whereas Gsdmd showed no significant change; HLWDD-M significantly reduced Nlrp3 and Il18 (*P* < 0.05) (Figure 3A–M). These results indicate that HLWDD attenuates NLRP3 inflammasome-related signaling in MS rat liver.

**Fig 3.**
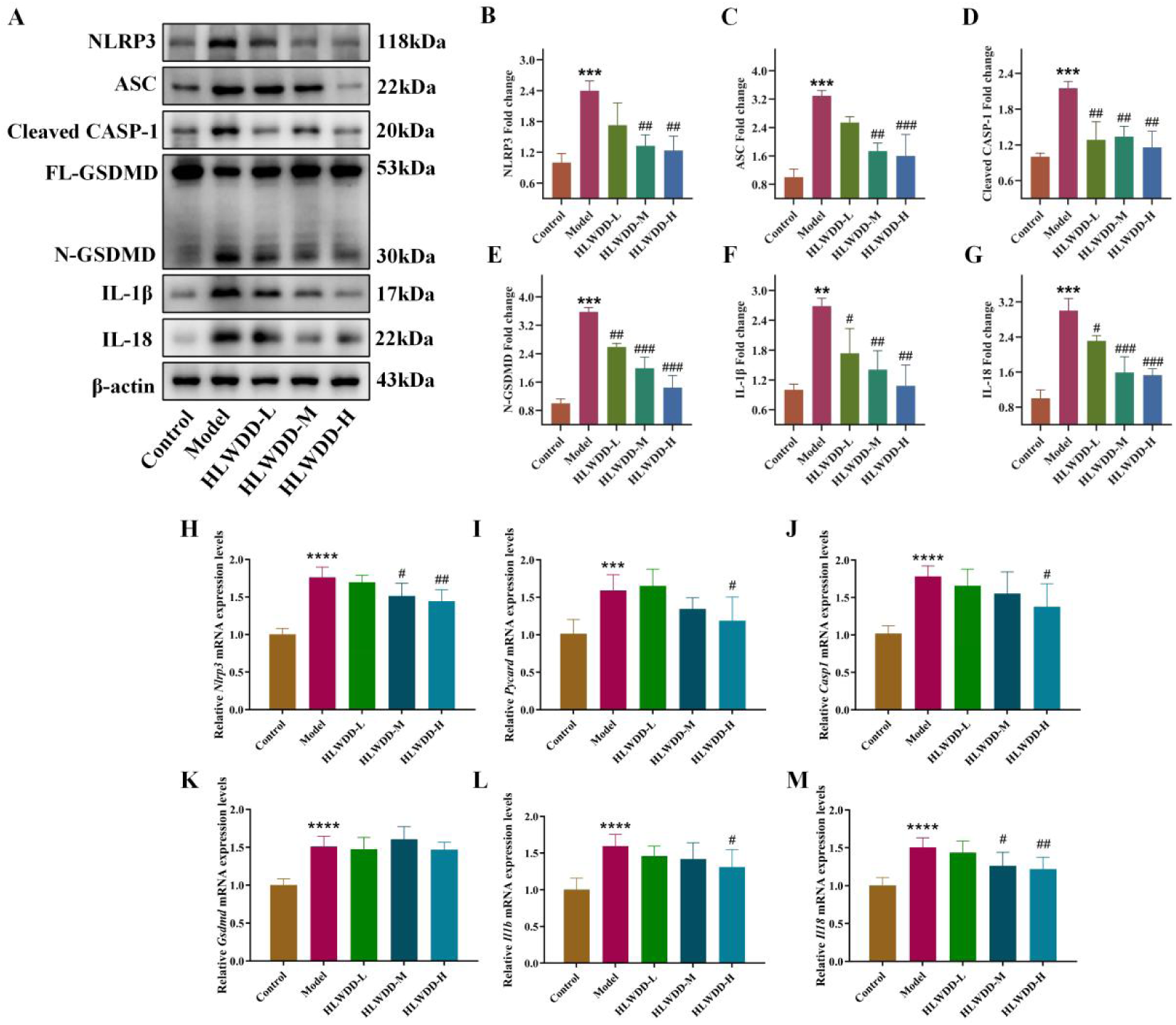
HLWDD attenuates NLRP3 inflammasome-related signaling in the liver of MS rats. (A) Representative Western blot images of NLRP3, ASC, cleaved CASP1, FL-GSDMD, N-GSDMD, IL-1β, and IL-18; (B–G) quantitative analysis of NLRP3 (B), ASC (C), cleaved CASP1 (D), N-GSDMD (E), IL-1β (F), and IL-18 (G); (H–M) qRT-PCR analysis of *Nlrp3* (H), *Pycard* (I), *Casp1* (J), *Gsdmd* (K), *Il1b* (L), and *Il18* (M). Data are expressed as mean ± SEM (n = 3 biological replicates). \*\**P*<0.01, \*\*\**P*<0.001, \*\*\*\**P*<0.0001, versus the control group; #*P*<0.05, ##*P*<0.01, ###*P*<0.001, ####*P*<0.0001, versus the model group.

#### Multi-Omics Integration: Gut–Liver Axis and NLRP3 Inhibition

Integrated analysis of the untargeted metabolomic profiles of colonic contents and liver tissues identified 117 and 67 differential metabolites, respectively, of which nine were shared between the two metabolic compartments. These shared metabolites included butyrate, valine, tryptophan, tyrosine, phosphatidylcholine, succinate, citrate, taurine, and glutathione, representing short-chain fatty acid, amino acid, phospholipid, organic acid, and sulfur-containing metabolic pathways (Figure 4A).

**Fig 4.**
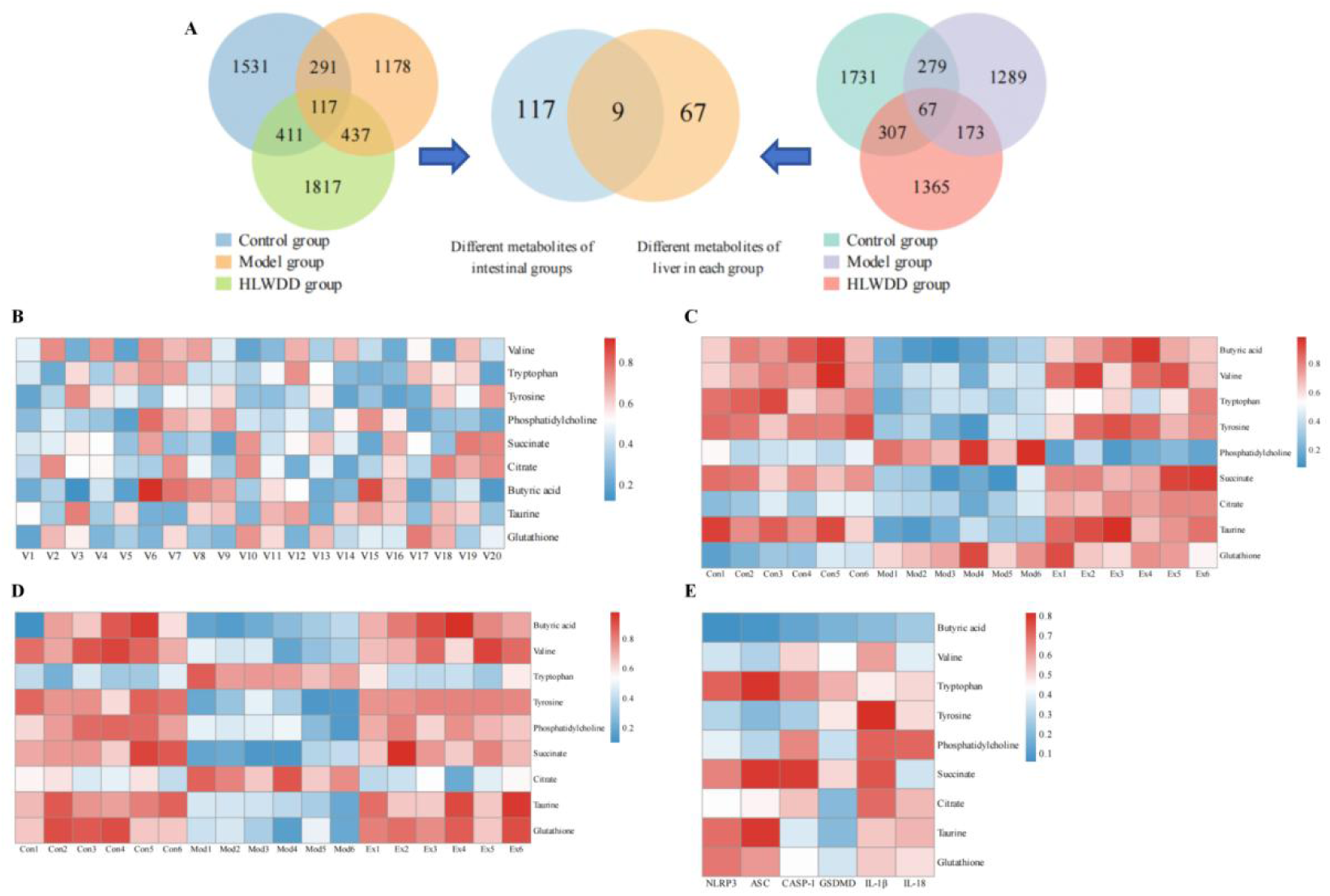
Integrated metagenomic and metabolomic analysis of HLWDD-mediated gut–liver metabolic regulation in MS. (A) Identification of shared differential metabolites between colonic contents and liver tissues. Venn diagrams show differential metabolites identified among the control, model, and HLWDD groups in colonic contents and liver tissues, respectively; nine metabolites were shared between the intestinal and hepatic metabolomic profiles. (B) Correlation heatmap between gut microbial features and the nine shared metabolites, including butyrate, valine, tryptophan, tyrosine, phosphatidylcholine, succinate, citrate, taurine, and glutathione. (C) Heatmap showing the relative abundance patterns of the nine shared metabolites in colonic contents from the control, model, and HLWDD groups. (D) Heatmap showing the relative abundance patterns of the nine shared metabolites in liver tissues from the control, model, and HLWDD groups. (E) Correlation heatmap between the nine shared metabolites and NLRP3 inflammasome-related proteins, including NLRP3, ASC, CASP1, GSDMD, IL-1 β, and IL-18. Each column in panels C and D represents one biological sample (n = 6 per group). Color intensity represents normalized metabolite abundance in panels C and D and the strength of the corresponding associations in panels B and E.

To further explore the relationship between gut microbial alterations and these shared metabolites, metagenomic data were integrated with the metabolomic profiles. Correlation analysis revealed distinct associations between gut microbial features and the nine shared metabolites. Notably, *Faecalibacterium* was strongly positively correlated with butyrate (r = 0.87), supporting a close association between this butyrate-producing genus and host butyrate metabolism (Figure 4B).

Heatmap analysis further demonstrated marked alterations in the abundance patterns of the nine shared metabolites in both colonic contents and liver tissues. Compared with the control group, the model group exhibited a substantially altered metabolic profile, whereas HLWDD treatment shifted the metabolite pattern toward that observed in the control group (Figure 4C,D). Butyrate was among the metabolites showing an apparent restoration following HLWDD intervention, consistent with the remodeling of gut – liver metabolic homeostasis.

Finally, correlation analysis between the nine shared metabolites and NLRP3 inflammasome-related proteins revealed extensive metabolite–protein associations involving NLRP3, ASC, CASP1, GSDMD, IL-1β, and IL-18 (Figure 4E). In particular, butyrate showed a strong association with multiple components of the NLRP3 inflammasome pathway. Collectively, these findings indicate that HLWDD-induced remodeling of the gut microbiota and gut – liver metabolic profile is closely associated with attenuation of NLRP3 inflammasome signaling, with the *Faecalibacterium* – butyrate relationship representing a potential link between microbial and inflammatory regulation.

#### Identification of Absorbed Components

UPLC-HRMS/MS analysis of HLWDD-containing rat serum yielded total ion chromatograms (Figure 5A–D). By searching against the ChemSpider and MassBank databases, a total of 32 chemical constituents were identified (S1 Table). These included ten organic acids and phenolic acids (1, 3, 4, 5, 7, 9, 10, 27, 29, 30), four flavonoids and chalcones (6, 8, 11, 12), two coumarins (14, 15), four alkaloids (13, 18, 20, 32), six terpenoids (16, 17, 19, 25, 28, 31), two steroids/triterpenoids (24, 26), one cyanogenic glycoside (2), and three other miscellaneous compounds (21, 22, 23).

**Fig 5.**
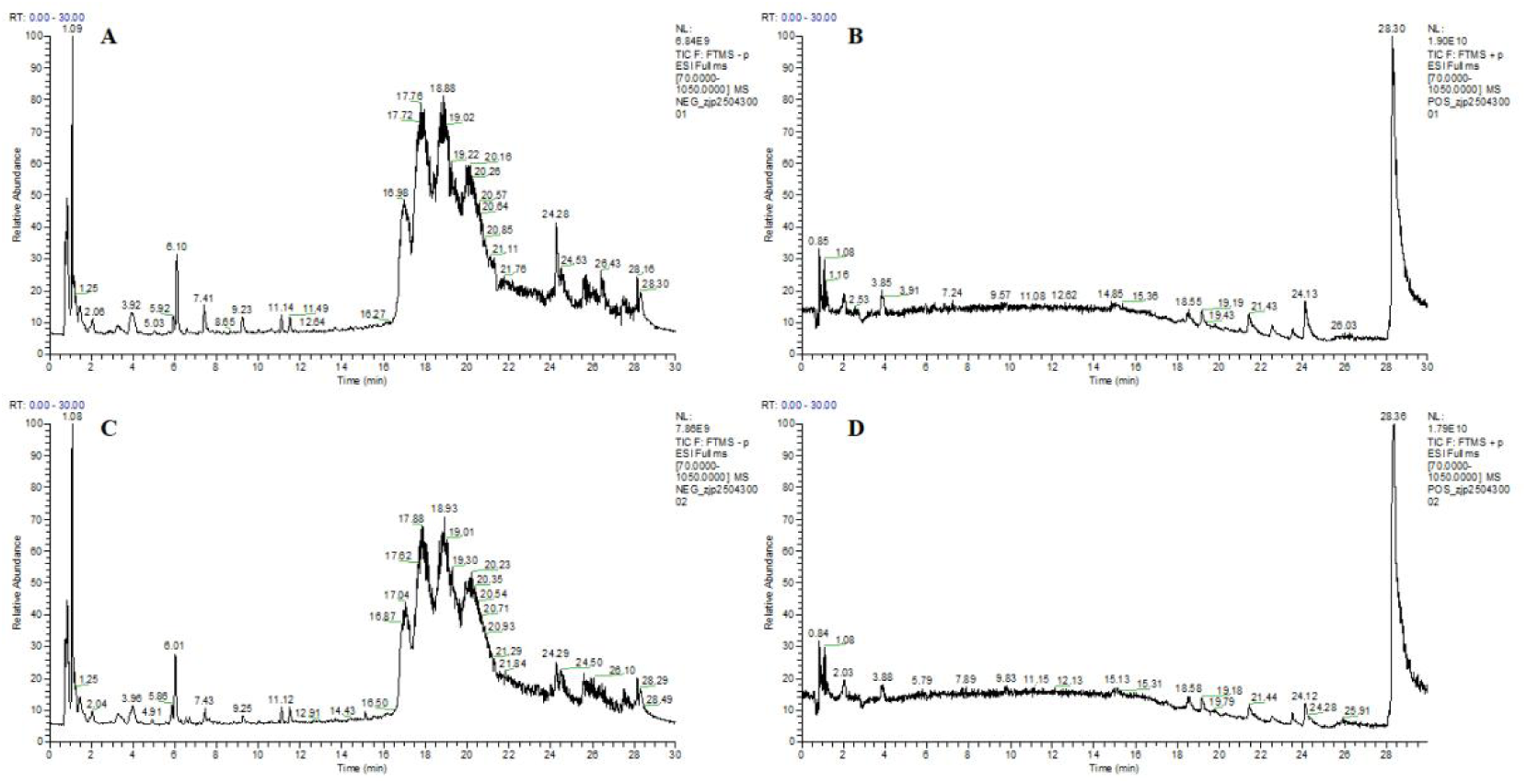
Chemical profiling of HLWDD-containing serum by UPLC-HRMS/MS. (A) Base peak chromatogram (BPC) of blank control serum in negative ion mode; (B) BPC of blank control serum in positive ion mode; (C) BPC of HLWDD-containing serum in negative ion mode; (D) BPC of HLWDD-containing serum in positive ion mode.

#### Network Pharmacology Analysis

Network pharmacology analysis identified 578 overlapping targets between HLWDD components and MS-related genes (Figure 6A). Thirty core targets were used to construct a protein – protein interaction (PPI) network, in which node size and color intensity reflected degree values. The top ten targets were AKT1, MAPK3, STAT3, CASP1, JUN, PIK3CA, SRC, MAPK1, PIK3R1, and ESR1 (Figure 6B). Forty-one candidate bioactive components were associated with these core targets.

**Fig 6.**
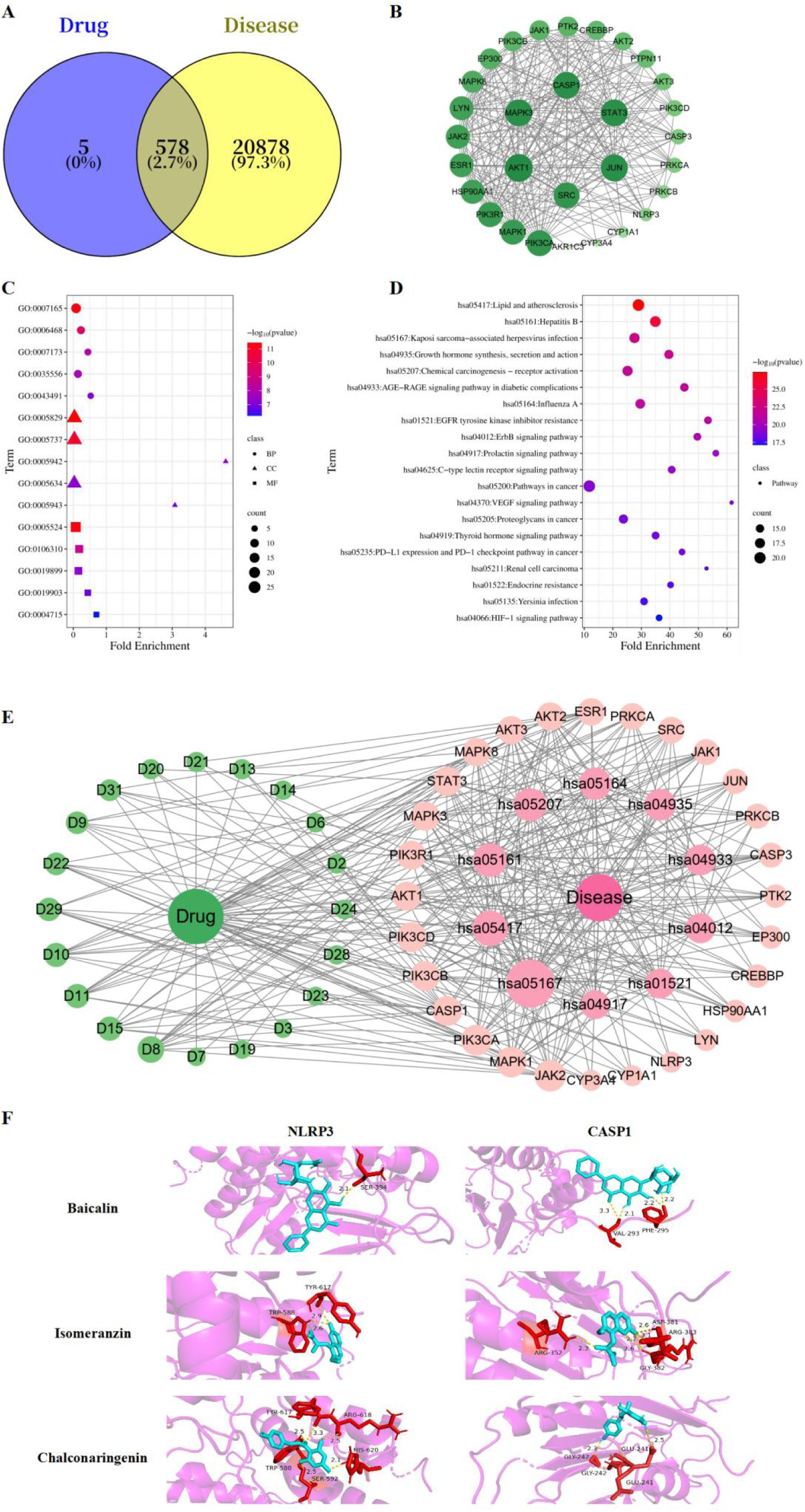
Network pharmacology and molecular docking analyses of HLWDD against MS. (A) Venn diagram showing overlapping targets between HLWDD components and MS-related genes; (B) PPI network of core targets; (C) GO enrichment analysis, including biological process, cellular component, and molecular function terms; (D) top 20 KEGG-enriched pathways; (E) compound–target– pathway network; (F) representative molecular docking poses of selected compounds with NLRP3 and CASP1. In network diagrams, node size and color intensity represent degree values; in enrichment plots, bubble color and size represent enrichment significance and gene count, respectively.

GO and KEGG enrichment analyses were performed using DAVID 6.8. A total of 391 GO terms were obtained, including 246 biological process (BP), 37 cellular component (CC), and 108 molecular function (MF) terms. BP terms were mainly associated with signal transduction, protein phosphorylation, epidermal growth factor receptor signaling, and PI3K/AKT pathway regulation (Figure 6C). CC terms were primarily related to the cytoplasm, cytosol, PI3K complex, and nucleus, whereas MF terms mainly involved ATP binding, serine/threonine kinase activity, enzyme binding, and phosphatase binding.

KEGG analysis identified 150 significantly enriched pathways, with the top 20 shown in Figure 6D. Representative pathways included lipid and atherosclerosis, AGE – RAGE signaling in diabetic complications, growth hormone synthesis, secretion and action, and chemical carcinogenesis – receptor activation. These pathways were mainly associated with lipid metabolism, oxidative stress, vascular regulation, and inflammatory responses relevant to MS.

A compound–target–pathway network was subsequently constructed using 20 representative compounds, 28 key targets, and 10 major KEGG pathways (Figure 6E). Baicalin, ferulic acid, chalconaringenin, eupalinolide O, and caffeic acid showed relatively high connectivity, while JAK2, MAPK1, PIK3CA, CASP1, PIK3CB, PIK3CD, MAPK3, AKT1, PIK3R1, and STAT3 were among the highly connected targets.

Molecular docking was further performed for baicalin, isomeranzin, and chalconaringenin with NLRP3 and CASP1. Four of the six ligand – protein combinations showed predicted binding energies below − 5.0 kcal/mol (Table 1, Figure 6F). Among the tested compounds, baicalin exhibited the most favorable predicted interactions with NLRP3 (−5.82 kcal/mol) and CASP1 (−5.41 kcal/mol), supporting its selection for subsequent experimental evaluation.

**Table 1.** Predicted binding energies of representative HLWDD compounds with NLRP3 and CASP1.

| NO. | Compound | Binding Energy /kcal·mol<sup>-1</sup> |  |
| --- | --- | --- | --- |
|  |  | NLRP3 | CASP1 |
| 1 | Baicalin | -5.82 | -5.41 |
| 2 | Isomeranzin | -5.19 | -5.03 |
| 3 | Chalconaringenin | -4.09 | -4.62 |

#### Quantification of Baicalin in HLWDD-Containing Rat Serum

Baicalin showed good linearity over the concentration range of 6.10–6,250.00 ng/mL. Weighted linear regression (1/x) yielded the calibration equation y = 170.11008x − 251.62241, with a correlation coefficient of r = 0.99983. The accuracy of the calibration standards ranged from 93.0% to 113.3% (Figure 7A). HPLC analysis of HLWDD-containing rat serum showed a baicalin concentration of 94.29 ng/mL (Figure 7B,C).

**Fig 7.**
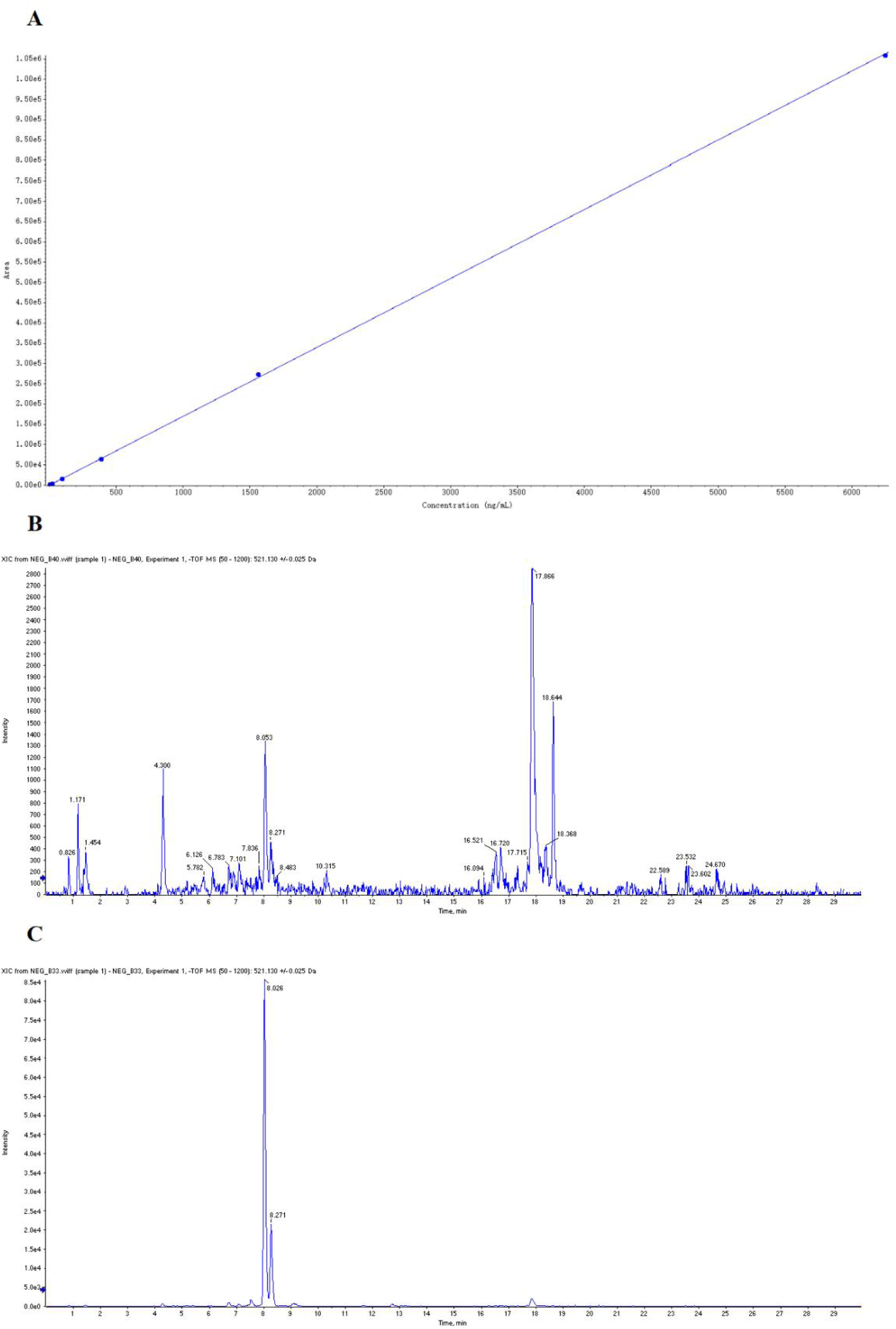
HPLC quantification of baicalin in HLWDD-containing rat serum. (A) Calibration curve of baicalin over the range of 6.10–6,250.00 ng/mL; (B) HPLC chromatogram of blank rat serum; (C) HPLC chromatogram of HLWDD-containing rat serum.

### *In Vitro* Cell-Based Validation

#### Effects of Baicalin on PA-BSA-Induced Lipotoxicity and Cytokine Production in HepG2 Cells

The CCK-8 assay showed that 80 μ M baicalin (BC) significantly reduced HepG2 cell viability (*P* < 0.05). Therefore, 15, 30, and 60 μ M BC were selected as the low, medium, and high concentrations for subsequent experiments (Figure 8A). PA-BSA significantly reduced cell viability, whereas BC-M and BC-H attenuated this effect (*P* < 0.01) (Figure 8B).

**Fig 8.**
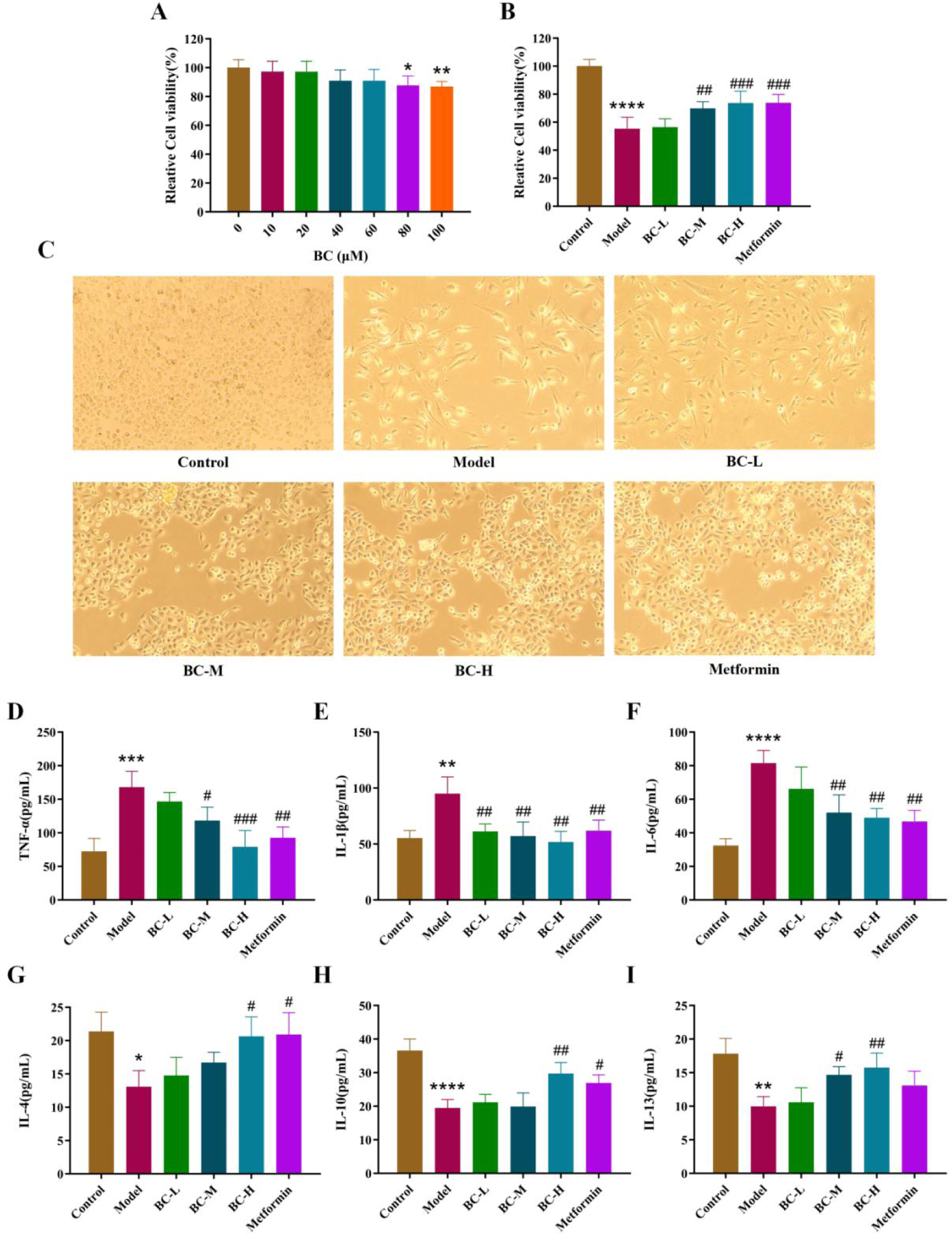
Baicalin attenuates PA-BSA-induced lipotoxicity and inflammatory responses in HepG2 cells. (A) Effects of different baicalin concentrations on HepG2 cell viability after 24 h, determined by CCK-8 assay; (B) effects of 15, 30, and 60 μM baicalin on PA-BSA-induced reduction in cell viability; (C) representative cell morphology observed under an inverted microscope; (D–I) levels of TNF-α (D), IL-1β (E), IL-6 (F), IL-4 (G), IL-10 (H), and IL-13 (I) in culture supernatants determined by ELISA. Data are expressed as mean ± SEM (n = 3 biological replicates). \**P*<0.05, \*\**P*<0.01, \*\*\**P*<0.001, \*\*\*\**P*<0.0001, versus the control group; #*P*<0.05, ##*P*<0.01, ###*P*<0.001, versus the model group.

Morphological examination showed that control cells exhibited a typical polygonal epithelial-like appearance and dense adherence, whereas PA-BSA-treated cells were reduced in number, shrunken, and spindle-shaped. Metformin treatment alleviated these morphological alterations. BC-L produced limited improvement, whereas BC-M partially restored cell morphology and BC-H resulted in cell density and morphology broadly comparable to those observed in the metformin group (Figure 8C).

ELISA showed that PA-BSA increased TNF-α, IL-1β, and IL-6 levels (*P* < 0.01) and decreased IL-4, IL-10, and IL-13 levels (*P* < 0.05) compared with the control group. BC-M and BC-H significantly reduced TNF-α, IL-1β, and IL-6 and increased IL-13 compared with the model group (*P* < 0.05), while BC-H additionally increased IL-4 and IL-10 (*P* < 0.05) (Figure 8D–I).

#### Effects of Baicalin on NLRP3 Inflammasome-Related Signaling in PA-BSA-Treated HepG2 Cells

Western blot analysis showed that PA-BSA significantly increased the protein levels of NLRP3, ASC, cleaved CASP1, N-GSDMD, IL-1 β, and IL-18 compared with the control group (*P* < 0.01), whereas BC-M and BC-H significantly reduced these protein levels (*P* < 0.05). At the mRNA level, BC-M significantly reduced NLRP3 and CASP1, whereas BC-H reduced NLRP3, PYCARD, CASP1, IL1B, and IL18 (*P* < 0.05); GSDMD showed no significant change (Figure 9A–M). These results indicate that baicalin attenuates NLRP3 inflammasome-related signaling in PA-BSA-treated HepG2 cells.

**Fig 9.**
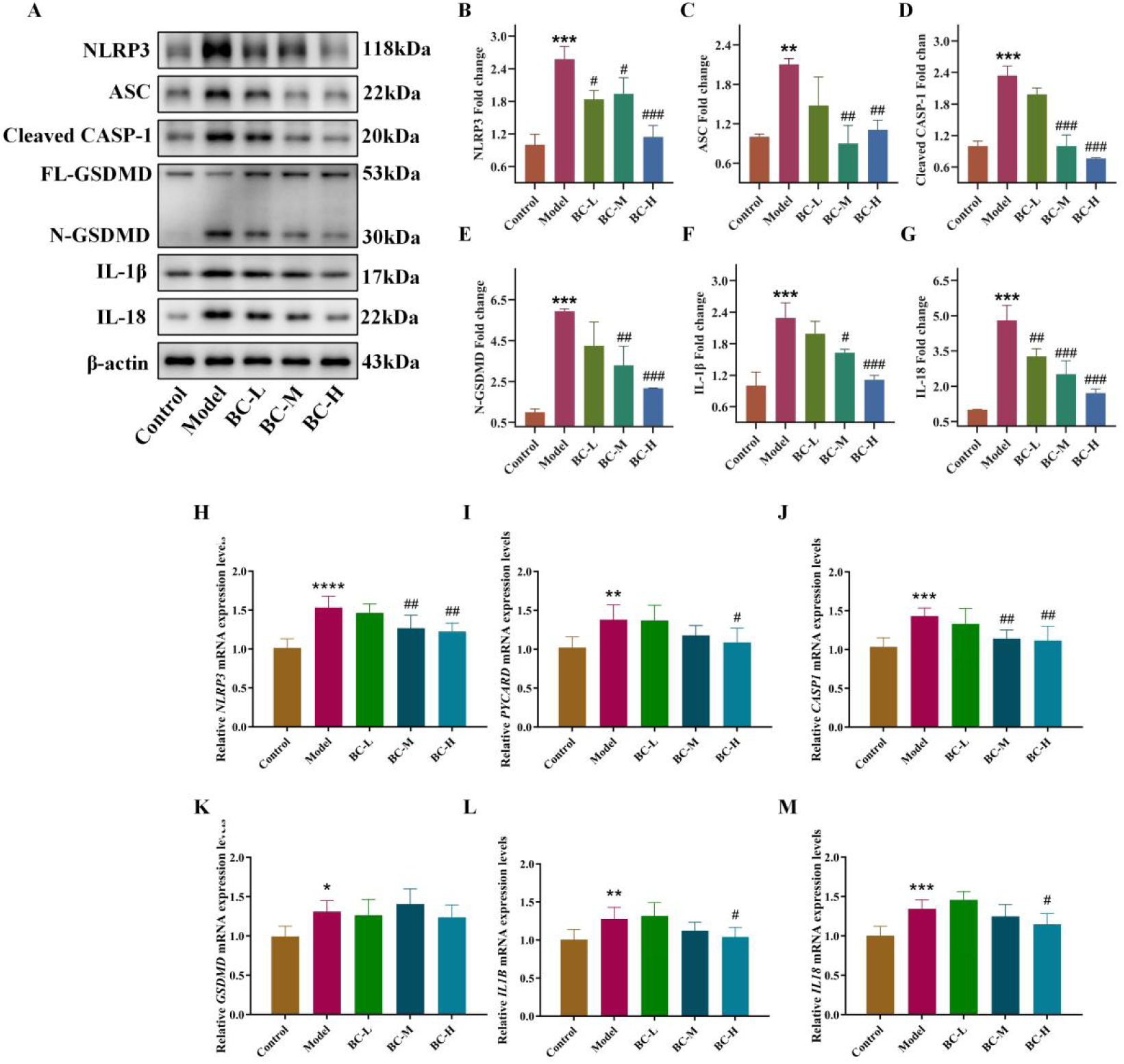
Baicalin attenuates NLRP3 inflammasome-related signaling in PA-BSA-treated HepG2 cells. (A) Representative Western blot images of NLRP3, ASC, cleaved CASP1, FL-GSDMD, N-GSDMD, IL-1β, and IL-18; (B–G) quantitative analysis of NLRP3 (B), ASC (C), cleaved CASP1 (D), N-GSDMD (E), IL-1β (F), and IL-18 (G); (H–M) qRT-PCR analysis of *NLRP3* (H), *PYCARD* (I), *CASP1* (J), *GSDMD* (K), *IL1B* (L), and *IL18* (M). Data are expressed as mean ± SEM (n = 3 biological replicates). \*\**P*<0.01, \*\*\**P*<0.001, \*\*\*\**P*<0.0001, versus the control group; #*P*<0.05, ##*P*<0.01, ###*P*<0.001, ####*P*<0.0001, versus the model group.

#### Targeted Metabolomic Analysis of Short-Chain Fatty Acids in PA-BSA-Treated HepG2 Cells

Compared with the control group, PA-BSA treatment significantly decreased acetate and butyrate levels and increased valerate and caproate levels (*P* < 0.05), whereas propionate remained unchanged. Compared with the model group, BC-H significantly increased the intracellular butyrate level (*P* < 0.05), while no significant changes were observed in the other SCFAs (Figure 10).

**Fig 10.**
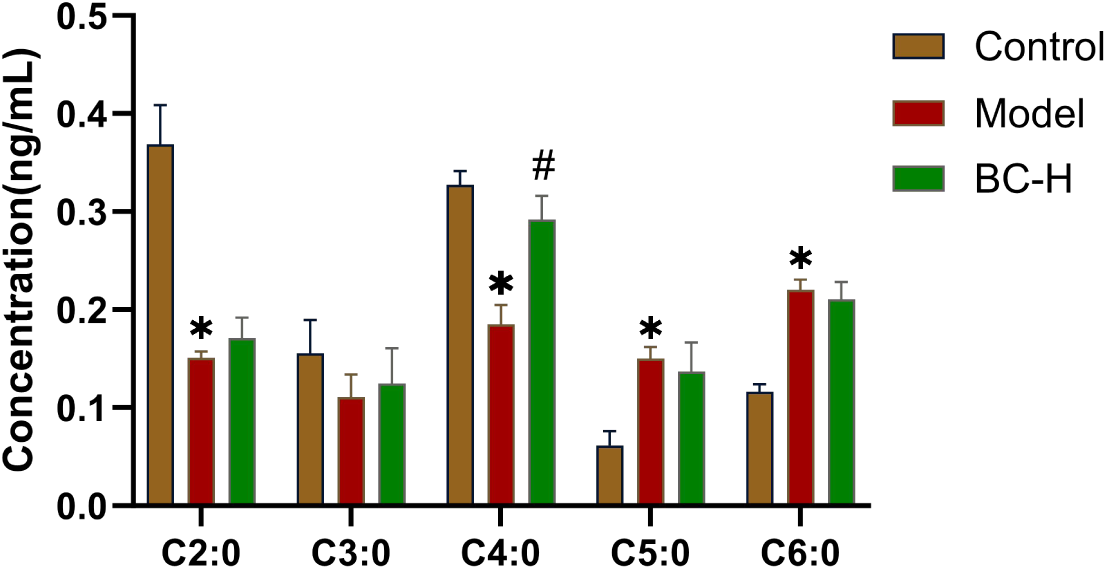
Effects of baicalin on intracellular short-chain fatty acid levels in PA-BSA-treated HepG2 cells. Intracellular levels of acetate (C2:0), propionate (C3:0), butyrate (C4:0), valerate (C5:0), and caproate (C6:0) were determined by targeted metabolomic analysis. Data are expressed as mean ± SEM (n = 6). \**P*<0.05, versus the control group; #*P*<0.05, versus the model group.

## Discussion

Chronic low-grade inflammation and metabolic dysregulation are central features of MS [23,24]. NLRP3 inflammasome activation promotes caspase-1-dependent maturation of IL-1β and IL-18 and contributes to insulin resistance and hepatic lipid accumulation [25]. Gut microbiota dysbiosis may further aggravate MS by disrupting microbial metabolite profiles, particularly SCFAs [26]. These observations support investigation of interventions that influence both metabolic and inflammatory pathways.

In the present study, HLWDD was evaluated as a defined multi-herbal pharmacological intervention in an experimentally induced MS model. Model establishment, treatment allocation, and outcome assessment were based on objective metabolic, biochemical, histological, molecular, and multi-omics measurements, without the use of traditional Chinese medicine syndrome or pattern classification. HLWDD improved glucose and lipid metabolism, hepatic steatosis, and systemic inflammation. Transcriptomic, Western blot, and qRT-PCR analyses consistently showed attenuation of NLRP3 inflammasome-related signaling, supporting an association between this pathway and the pharmacological effects of HLWDD.

Integrated analysis of colonic and hepatic metabolomic profiles identified nine shared differential metabolites. Metagenomic – metabolomic correlation analysis further revealed a strong positive association between *Faecalibacterium* and butyrate. As *Faecalibacterium* is an important butyrate-producing commensal, these findings suggest that changes in butyrate-related microbial metabolism may accompany the metabolic effects of HLWDD. Butyrate was also associated with several NLRP3 inflammasome-related proteins. However, these correlations do not establish a causal *Faecalibacterium* – butyrate – NLRP3 pathway and require targeted microbiota and metabolite intervention studies for confirmation.

UPLC-HRMS/MS analysis tentatively identified 32 serum-absorbed constituents of HLWDD. Network pharmacology and molecular docking indicated that baicalin showed relatively high network connectivity and favorable predicted interactions with NLRP3 and CASP1. HPLC further confirmed baicalin in HLWDD-containing serum at 94.29 ng/mL. Together with previous evidence that baicalin can inhibit NLRP3 inflammasome-related signaling [30], these findings support baicalin as a candidate absorbed bioactive constituent rather than establishing it as the principal active component.

In PA-BSA-treated HepG2 cells, baicalin attenuated lipotoxicity, improved inflammatory cytokine profiles, and reduced NLRP3 inflammasome-related signaling. The highest tested concentration also increased intracellular butyrate. However, the *in vitro* concentrations exceeded the measured circulating baicalin concentration, and HepG2 cells lack a gut microbial component. These findings should therefore be interpreted as supportive cellular evidence rather than a direct simulation of *in vivo* exposure or microbiota-derived butyrate production.

Overall, HLWDD ameliorated metabolic and inflammatory abnormalities in experimental MS and was associated with restoration of gut–liver metabolic homeostasis and attenuation of NLRP3 inflammasome-related signaling. Further microbiota intervention, metabolite rescue, pharmacokinetic, and direct target-validation studies are required to establish causality.

## Supporting information

S1 Table. Chemical constituents identified in HLWDD by UPLC-HRMS/MS. Retention times, mass spectral data, and compound classifications.

## Abbreviations

ASC: Apoptosis-Associated Speck-Like Protein Containing a CARD
BCA: Bicinchoninic Acid
CASP1: Caspase-1
CCK-8: Cell Counting Kit-8
ECL: Enhanced Chemiluminescence
FBG: Fasting Blood Glucose
GC-MS: Gas Chromatography-Mass Spectrometry
GO: Gene Ontology
GSDMD: Gasdermin D
HDL-C: High-Density Lipoprotein Cholesterol
H&E: Hematoxylin and Eosin
HPLC: High-Performance Liquid Chromatography
HLWDD: Huanglian Wendan Decoction
HRMS: High-Resolution Mass Spectrometry
IL: Interleukin
KEGG: Kyoto Encyclopedia of Genes and Genomes
LDL-C: Low-Density Lipoprotein Cholesterol
MS: Metabolic Syndrome
NLRP3: NLR Family Pyrin Domain Containing 3
PA-BSA: Palmitic Acid-Bovine Serum Albumin
PPI: Protein-Protein Interaction
PVDF: Polyvinylidene Fluoride
qRT-PCR: Quantitative Real-Time Polymerase Chain Reaction
SCFAs: Short-Chain Fatty Acids
TC: Total Cholesterol
TCM: Traditional Chinese Medicine
TG: Triglycerides
TNF-α: Tumor Necrosis Factor-α
UPLC-HRMS/MS: Ultra-Performance Liquid Chromatography-High-Resolution Tandem Mass Spectrometry

## Institutional Review Board Statement

All animal experimental procedures were approved by the Animal Ethics Committee of Hunan University of Chinese Medicine (LLBH-LL2025121708) and conducted in accordance with institutional guidelines for the care and use of laboratory animals.

## Informed Consent Statement

Not applicable.

## Data Availability Statement

The data supporting the findings of this study are available from the corresponding author upon reasonable request.

## Acknowledgments

Not applicable.

## References

1. Neeland IJ, Lim S, Tchernof A, Gastaldelli A, Rangaswami J, Ndumele CE, et al. Metabolic syndrome. Nat Rev Dis Primers. 2024;10:77. 10.1038/s41572-024-00563-5

2. Powell EE, Wong VW, Rinella M. Non-alcoholic fatty liver disease. Lancet. 2021;397:2212–2224. 10.1016/S0140-6736(20)32511-3

3. Rohm TV, Meier DT, Olefsky JM, Donath MY. Inflammation in obesity, diabetes, and related disorders. Immunity. 2022;55:31–55. 10.1016/j.immuni.2021.12.013

4. Li S, Zhang T, Yang H, Chang Q, Zhao Y, Chen L, et al. Metabolic syndrome, genetic susceptibility, and risk of chronic obstructive pulmonary disease: The UK Biobank Study. Diabetes Obes Metab. 2024;26:482–494. 10.1111/dom.15334

5. Petrenko V, Sinturel F, Riezman H, Dibner C. Lipid metabolism around the body clocks. Prog Lipid Res. 2023;91:101235. 10.1016/j.plipres.2023.101235

6. Sandsdal RM, Juhl CR, Jensen SBK, Lundgren JR, Janus C, Blond MB, et al. Combination of exercise and GLP-1 receptor agonist treatment reduces severity of metabolic syndrome, abdominal obesity, and inflammation: a randomized controlled trial. Cardiovasc Diabetol. 2023;22:41. 10.1186/s12933-023-01765-z

7. Fu J, Wu H. Structural mechanisms of NLRP3 inflammasome assembly and activation. Annu Rev Immunol. 2023;41:301–316. 10.1146/annurev-immunol-081022-021207

8. Huang Y, Xu W, Zhou R. NLRP3 inflammasome activation and cell death. Cell Mol Immunol. 2021;18:2114–2127. 10.1038/s41423-021-00740-6

9. Wan X, Xu C, Lin Y, Lu C, Li D, Sang J, et al. Uric acid regulates hepatic steatosis and insulin resistance through the NLRP3 inflammasome-dependent mechanism. J Hepatol. 2016;64:925–932. 10.1016/j.jhep.2015.11.022

10. Sharma BR, Kanneganti TD. NLRP3 inflammasome in cancer and metabolic diseases. Nat Immunol. 2021;22:550–559. 10.1038/s41590-021-00886-5

11. Boulangé CL, Neves AL, Chilloux J, Nicholson JK, Dumas ME. Impact of the gut microbiota on inflammation, obesity, and metabolic disease. Genome Med. 2016;8:42. 10.1186/s13073-016-0303-2

12. Zeng SL, Li SZ, Xiao PT, Cai YY, Chu C, Chen BZ, et al. Citrus polymethoxyflavones attenuate metabolic syndrome by regulating gut microbiome and amino acid metabolism. Sci Adv. 2020;6:eaax6208. 10.1126/sciadv.aax6208

13. Michels N, Zouiouich S, Vanderbauwhede B, Vanacker J, Indave Ruiz BI, Huybrechts I. Human microbiome and metabolic health: an overview of systematic reviews. Obes Rev. 2022;23:e13409. 10.1111/obr.13409

14. Chen B, Guan L, Wu C, Gong Y, Wu L, Zhang M, et al. Gut microbiota-butyrate-PPARγ axis modulates adipose regulatory T cell population. Adv Sci (Weinh). 2025;12:e2411086. 10.1002/advs.202411086

15. Bridgeman SC, Northrop W, Melton PE, Ellison GC, Newsholme P, Mamotte CDS. Butyrate generated by gut microbiota and its therapeutic role in metabolic syndrome. Pharmacol Res. 2020;160:105174. 10.1016/j.phrs.2020.105174

16. Liu ZC, Fu HJ, Li NC, Deng FJ, Gan YK, Ye YJ, et al. Systematic pharmacology and experimental validation to elucidate the inflammation-associated mechanism of Huanglian Wendan (HLWD) decoction in the treatment of MAFLD associated with atherosclerosis. J Ethnopharmacol. 2025;337:118841. 10.1016/j.jep.2024.118841

17. Li YB, Zhang WH, Liu HD, Liu Z, Ma SP. Protective effects of Huanglian Wendan Decoction against cognitive deficits and neuronal damages in rats with diabetic encephalopathy by inhibiting the release of inflammatory cytokines and repairing insulin signaling pathway in hippocampus. Chin J Nat Med. 2016;14:813–822. 10.1016/S1875-5364(16)30098-X

18. Su Z, Li Y, Zhou Z, Tang L, Zhu Y, Chen H, et al. Huanglian Wendan decoction alleviates metabolic syndrome by reprogramming macrophage polarization: evidence from multi-omics integration. Phytomedicine. 2025;146:157119. 10.1016/j.phymed.2025.157119

19. Zhu J, Chen Y, Han Y, Li J. Mechanism of Huanglian Wendan Decoction in ameliorating non-alcoholic fatty liver disease via modulating gut microbiota-mediated metabolic reprogramming and activating the LKB1/AMPK pathway. PLoS One. 2025;20:e0331303. 10.1371/journal.pone.0331303

20. Jia KK, Ding H, Yu HW, Dong TJ, Pan Y, Kong LD. Huanglian-Wendan decoction inhibits NF-κB/NLRP3 inflammasome activation in liver and brain of rats exposed to chronic unpredictable mild stress. Mediators Inflamm. 2018;2018:3093516. 10.1155/2018/3093516

21. He Y, Wang H, Lin S, Chen T, Chang D, Sun Y, et al. Advanced effect of curcumin and resveratrol on mitigating hepatic steatosis in metabolic associated fatty liver disease via the PI3K/AKT/mTOR and HIF-1/VEGF cascade. Biomed Pharmacother. 2023;165:115279. 10.1016/j.biopha.2023.115279

22. Geng Y, Hernández Villanueva A, Oun A, Buist-Homan M, Blokzijl H, Faber KN, et al. Protective effect of metformin against palmitate-induced hepatic cell death. Biochim Biophys Acta Mol Basis Dis. 2020;1866:165621. 10.1016/j.bbadis.2019.165621

23. Islam MS, Wei P, Suzauddula M, Nime I, Feroz F, Acharjee M, et al. The interplay of factors in metabolic syndrome: understanding its roots and complexity. Mol Med. 2024;30:279. 10.1186/s10020-024-01019-y

24. Kawai T, Autieri MV, Scalia R. Adipose tissue inflammation and metabolic dysfunction in obesity. Am J Physiol Cell Physiol. 2021;320:C375–C391. 10.1152/ajpcell.00379.2020

25. Liu D, Zhou C, Wang X, Luo Z, Xu R, Huo S, et al. Nucleotide metabolic rewiring enables NLRP3 inflammasome hyperactivation in obesity. Science. 2026;391:eadq9006. 10.1126/science.adq9006

26. Liu W, Yu L, Chen Q, Zhang C, Wang L, Yu N, et al. Poria cocos polysaccharides alleviate obesity-related adipose tissue insulin resistance via gut microbiota-derived short-chain fatty acids activation of FGF21/PI3K/AKT signaling. Food Res Int. 2025;215:116671. 10.1016/j.foodres.2025.116671

27. Shi Z, Li M, Zhang C, Li H, Zhang Y, Zhang L, et al. Butyrate-producing Faecalibacterium prausnitzii suppresses natural killer/T-cell lymphoma by dampening the JAK-STAT pathway. Gut. 2025;74:557–570. 10.1136/gutjnl-2024-333530

28. Yu T, Cao A, Zhu F, Xie Z, Hu S, Li D. Butyrate ameliorates ISO-induced cardiac and intestinal injury in rats via modulation of bitter taste receptors (Tas2rs) and GPR41/43 to inhibit NLRP3 activation. Nutrients. 2026;18:1530. 10.3390/nu18101530

29. Zhao Z, Tong Y, Kang Y, Qiu Z, Li Q, Xu C, et al. Sodium butyrate (SB) ameliorated inflammation of COPD induced by cigarette smoke through activating the GPR43 to inhibit NF-κB/MAPKs signaling pathways. Mol Immunol. 2023;163:224–234. 10.1016/j.molimm.2023.10.007

30. Zhao J, Wang Z, Yuan Z, Lv S, Su Q. Baicalin ameliorates atherosclerosis by inhibiting NLRP3 inflammasome in apolipoprotein E-deficient mice. Diab Vasc Dis Res. 2020;17:1479164120977441. 10.1177/1479164120977441

31. Bonnet F, Scheen A. Understanding and overcoming metformin gastrointestinal intolerance. Diabetes Obes Metab. 2017;19:473–481. 10.1111/dom.12854

